# Resolving within-host malaria parasite diversity using single-cell sequencing

**DOI:** 10.1101/391268

**Authors:** Standwell C. Nkhoma, Simon G. Trevino, Karla M. Gorena, Shalini Nair, Stanley Khoswe, Catherine Jett, Roy Garcia, Benjamin Daniel, Aliou Dia, Dianne J. Terlouw, Stephen A. Ward, Timothy J.C. Anderson, Ian H. Cheeseman

## Abstract

Malaria patients can carry one or more clonal lineage of the parasite, *Plasmodium falciparum*, but the composition of these infections cannot be directly inferred from bulk sequence data. Well-defined, complete haplotypes at single-cell resolution are ideal for describing within-host population structure and unambiguously determining parasite diversity, transmission dynamics and recent ancestry but have not been analyzed on a large scale. We generated 485 near-complete single-cell genome sequences isolated from fifteen *P. falciparum* patients from Chikhwawa, Malawi, an area of intense malaria transmission. Matched single-cell and bulk genomic analyses revealed patients harbored up to seventeen unique lineages. Estimation of parasite relatedness within patients suggests superinfection by repeated mosquito bites is rarer than co-transmission of parasites from a single mosquito. Our single-cell analysis indicates strong barriers to establishment of new infections in malaria-infected patients and allows high resolution dissection of intra-host variation in malaria parasites.

---

Within a single host interactions between genetically distinct malaria parasites influence the evolution of parasite virulence, antimalarial drug resistance, immunity, gametocyte sex ratios, and malaria transmission in mouse malaria models^1-5^. Complex infections (that contain more than one unique parasite genetic background) confound most traditional genetic analysis, preventing the accurate inference of allele frequencies and even simple phenotype associations^6,7^. Complex infections play a key role in the structure of populations as they provide the substrate for sexual recombination to occur, in turn shaping local decay of linkage disequilibrium and haplotype variation^8-10^. The impact of within-host interactions remain largely unknown in human malaria. This is due to the paucity of appropriate tools for resolving infection complexity on a large-scale at the level of single parasitized cells: we cannot directly infer the composition of malaria infections by bulk sequencing of infected blood samples. Important insights into the genetic architecture of individual malaria infections are emerging, aided by recent advances in targeted capture of singly-infected erythrocytes from complex mixtures and improved methods for single-cell sequencing^7,11^, and computational approaches for interpreting infection complexity^12,13^.

Genetically distinct malaria parasites can infect an individual through two routes (Fig. 1). A single individual may be bitten by two (or more) infected mosquitos, each bearing a unique parasite genotype, or an individual may be bitten by a single mosquito bearing more than one parasite genotype. Throughout, we refer to these two processes as superinfection and co-transmission respectively. Following a bloodmeal, gametocyte stage parasites fuse in the mosquito midgut, and an obligate round of sexual recombination occurs. If only a single parasite genotype is present all offspring will be identical (Fig. 1, top panel), with the presence of multiple genotypes allowing recombinant progeny to arise^6,8,14-16^ (Fig. 1, bottom panel). Using single cell sequencing and cloning by limiting dilution of parasites from a single individual we have previously seen a range of inferred relationships, including identical clones, siblings and unrelated individuals^7,11,16^. However, it is unknown to what extent these findings can be generalized across a population. The degree to which the genetic diversity of individual infections is driven by superinfection of unrelated strains, or co-transmission of related ones is needed to model how genetic diversity could be maintained in the face of malaria control measures.

**Figure 1.**
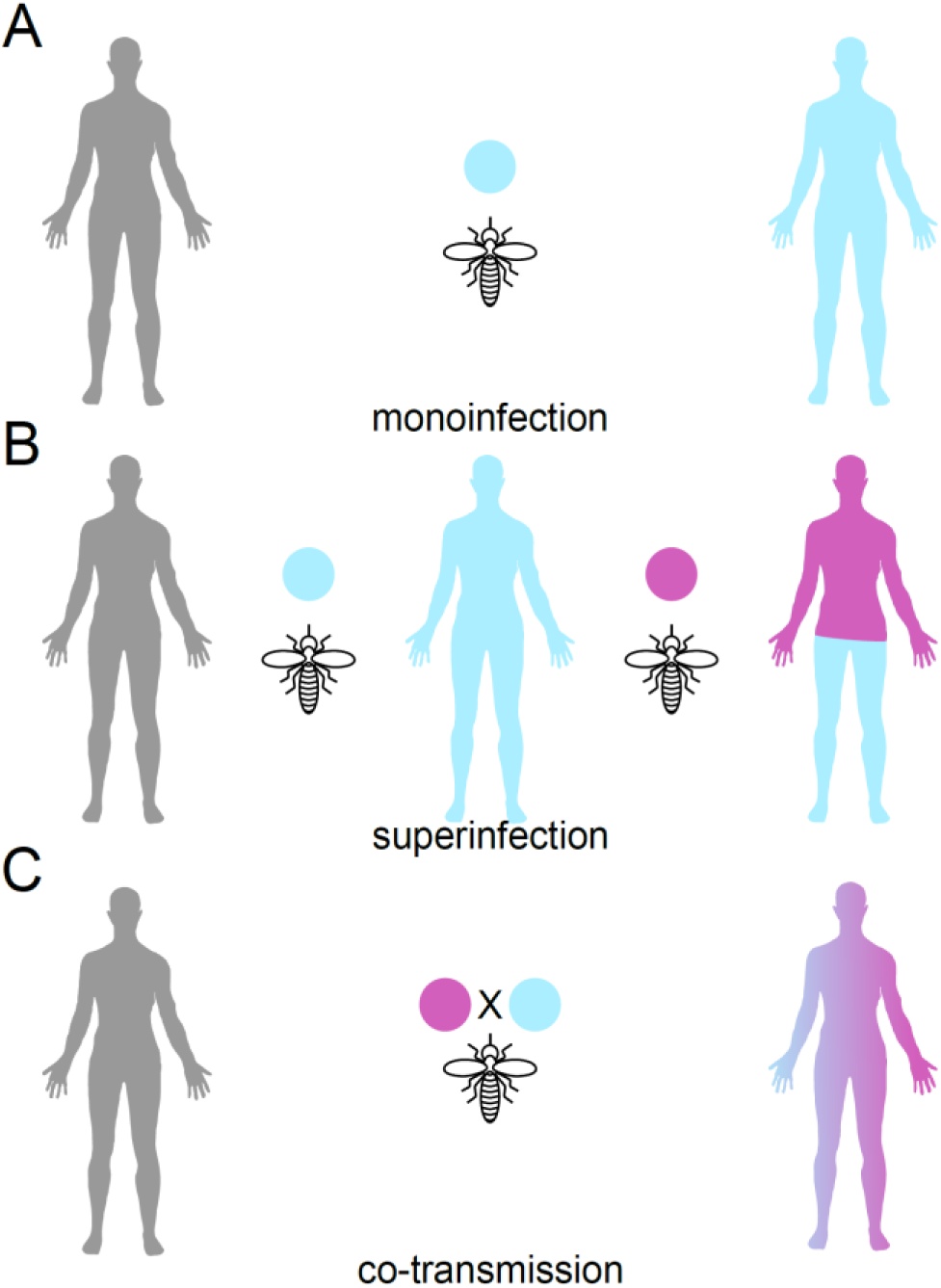
The within host genetic diversity of malaria parasites is shaped by transmission strategy. (A) A simple monoinfection is generated when an uninfected individual is bitten by a mosquito bearing a single parasite genotype. (B) A superinfection occurs when an individual is bitten by two mosquitos, each bearing a single parasite genotype. (C) Co-transmission of parasites occurs when a single mosquito bearing multiple genetically distinct parasites bites an uninfected individual. As genetic recombination is an obligate stage of mosquito transmission multiple related parasites may infected through this route.

## Infection complexity in bulk sequenced samples

To resolve the within-host structure of malaria infections, we performed a cross-sectional survey of individuals infected with uncomplicated *P. falciparum* malaria in Chikhwawa, Malawi, an area of high malaria transmission (entomological inoculation rate 183 infectious bites per person per year^17^). We performed bulk parasite genome sequencing of 49 infections to a median read depth of 31 (interquartile range 20.93-48.37). We estimated the complexity of infection of bulk sequence data using 10,997 unfixed SNP positions with a minor allele frequency (MAF) >0.05 using the F_WS_ statistic^18,19^ and DEploid^13^ (Fig. 2a,b, Supplementary Table 1). F_WS_ grades infections on a continuous scale of complexity where infections with an F_WS_>0.95 are considered clonal and DEploid estimates the number of haplotypes (K) present in sequence data by jointly estimating haplotypes and their abundances. In close agreement with contemporary estimates of within host diversity^20^, 22 of 49 infections (44.9%) were considered clonal by F_WS_. The within-host allele frequency (WHAF) captured from deep sequencing can be used to infer the presence of related parasites^21^. The patterns of unfixed mutations in the remaining 27 infections suggest a simple model of superinfection, where two unrelated parasite genetic backgrounds colonize an individual, are insufficient to universally capture all patterns of within-host relatedness (Supplementary File 1). We selected 15 infections across the range of F_WS_ and inferred K for single-cell sequencing, using a recently optimized method to generate near-complete genome capture^11^. The malaria parasite undergoes 4-5 rounds of DNA replication within a single cell producing segmented schizont stage parasites with an average of 16 genome copies^22^. We isolate individual schizonts by fluorescence activated cell sorting, followed by whole genome amplification (WGA) under highly sterile conditions before sequencing the amplified product.

**Figure 2.**
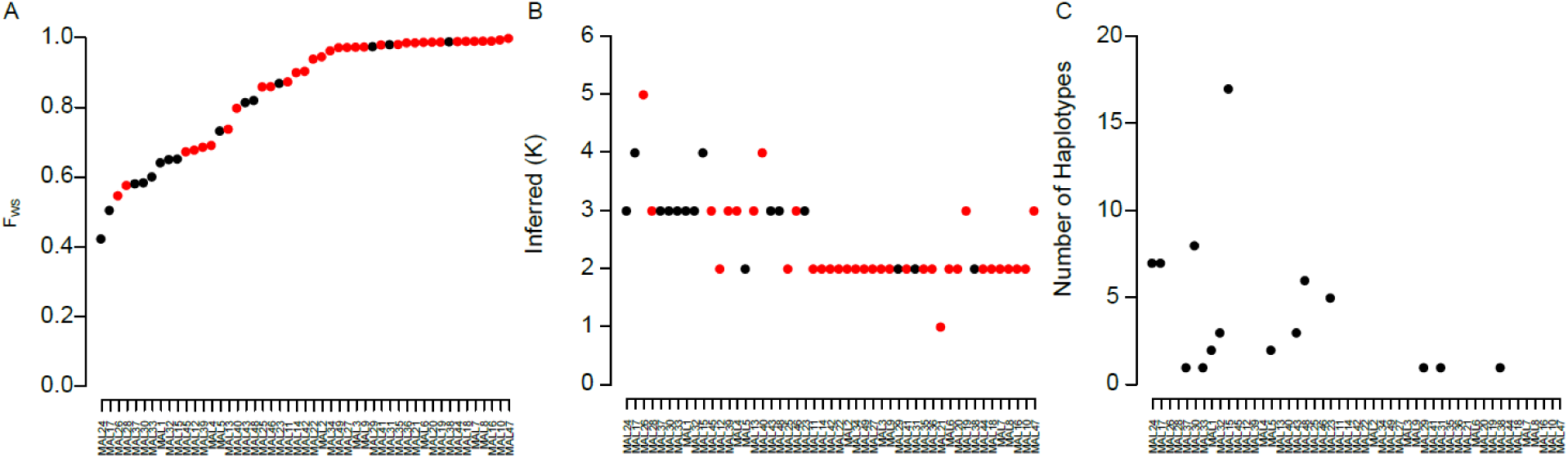
Complexity of infection inferred from bulk and single-cell sequencing. (A) F_WS_ scores for 49 bulk sequenced infections. Infections above the dashed line (F_WS_=0.95) are assumed to be clonal. (B) Inferred number of haplotypes (K) inferred by DEploid, infections are ordered by the F^WS^ score. Black dots in (A) and (B) denote infections also deconvoluted by single-cell sequencing. (C) Number of unique haplotypes inferred by single-cell sequencing.

## Single cell sequencing of malaria parasites

In total we sequenced the genomes of 485 single-cells subjected to WGA (437 unique to this study), 49 bulk infections and 24 clones isolated from a single patient by limiting dilution^16,23^. Prior to genotype filtering we scored 175,543 biallelic SNPs with a VQSLOD>0 across the 558 genome sequences. The highly repetitive and AT-rich *P. falciparum* genome^24^presents unique challenges with generating an accurate picture of the variation present in a single-cell. We were particularly concerned with capture of DNA from more than one genetic background during the single-cell sequencing protocol and implemented stringent quality checks. Using sequencing data from the 24 clones we estimated the threshold for identifying single cell sequences where there was potential contamination from exogenous DNA at 1% of mixed base calls. The sequences from the cloned lines were integrated into the single cell dataset for downstream analysis. After excluding low coverage libraries (<75,000 calls, n=23) and sequences with >1% mixed base calls (n=38) 424 single-cell sequences remained. After including 23 of the sequences from *ex vivo* expanded clones there were 13-45 sequences per infection (mean 29.9 sequences; Supplementary Data Fig. 1). The number of sequences per sample attempted was determined by rarefaction analysis (described below).

After quality control of the dataset we retained 60,002 SNPs scored in at least 90% of the 496 sequences, 10,997 of which had a MAF>0.05 across the 49 bulk sequenced infections. As an initial characterization of our data we estimated the genetic diversity in each infection from the number of unfixed sites from read pileups in bulk sequencing or across called genotypes in single-cell sequencing. For paired bulk/single-cell data from the same infection a mean of 1.6 fold (range 0.7-9.1 fold) more polymorphic sites were discovered by single-cell sequencing than by bulk sequencing (Supplementary Data Fig. 2). This is likely due to the limits in discovery of very low frequency SNPs by bulk sequencing. By subsampling our single-cell data we saw diminishing returns from sequencing additional cells, with 90% of the observed polymorphic sites captured by sampling a mean of 21.6 cells (range 7-43, Supplementary Data Fig. 2).

## Haplotypic diversity of malaria infections

A major goal in malaria genomics has been estimating the number of unique haplotypes (or complexity of infection) within an infection^25^. We estimated the number of unique haplotypes directly from the single-cell data. To exclude potential confounding of *de novo* mutation and sequencing error we restricted analysis to 10,997 conservatively called sites with a MAF >0.05 in the 49 bulk sequenced infections. We estimated the number of unique haplotypes per infection by collapsing haplotypes from the same infection that were different at <1% of sites. For each infection we applied individual-based rarefaction to the haplotype abundances and sequenced additional single genomes until a plateau in the rarefaction curve was reached (Supplementary Data Fig. 3). Using this approach between 1 and 17 haplotypes were observed in each infection (Fig. 1c, Supplementary Data Table 1). There was strong correlation between the effective number of strains^13^ inferred by single cell sequencing and the effective K from DEploid (Pearson’s *r*^2^=0.61) and F_WS_ (Pearson’s *r*^2^=-0.51, Supplementary Data Fig. 4). Rarefaction of haplotype abundance suggested exhaustive capture of haplotypes in 12/15 infections. In two infections (MAL23 and MAL30) we sampled two fewer haplotypes than suggested by rarefaction (Chao I estimator-MAL23=6.94, MAL30=10.18, observed haplotypes-MAL23=5, MAL30=8). In both cases the observed number of haplotypes were within the 95% confidence intervals of the estimation. One infection (MAL15) showed exceptionally high diversity with 17 of an estimated 30.21 (95% CI=19.7-81.7) haplotypes detected. Two infections (MAL37 and MAL33) show a single haplotype from single-cell sequencing, although F_WS_ scores <0.95 and patterns of segregating sites suggest we have incompletely captured all haplotypes (Supplementary File 1). Sequencing more cells did not capture additional haplotypes.

## Population structure of individual infections

The number of unique haplotypes alone captures only a single aspect of the recent history of genetically distinct parasites from the same infection. For instance it does not distinguish whether diversity is due to superinfection of unrelated strains from multiple mosquito inoculations, from co-transmission of related strains from a single mosquito inoculation or from a combination of the two. To jointly characterize the between- and within-host genetic structure of malaria infections we clustered infections based upon either a UPGMA tree of pairwise allele sharing (Fig. 3a) or by unsupervised clustering in ADMIXTURE^26^ (K=31; Fig. 3b, Supplementary Data Figure 5). In both cases sequences from the same infection predominantly cluster together, suggesting they were more related to each other than parasites drawn from the population and diversity is likely the result of co-transmission by related parasites from the same mosquito.

**Figure 3.**
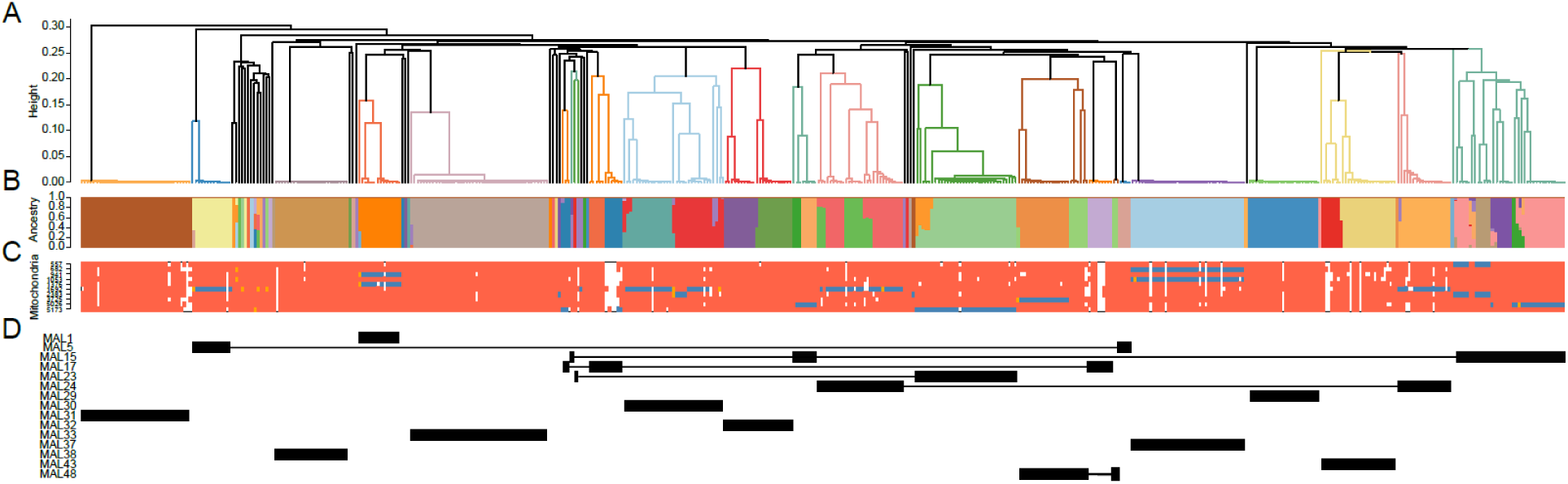
Clustering of single-cell and bulk genome sequences. (A) A UPGMA tree of 1-pairwise allele sharing across all samples passing quality control. Shading behind the tree and labels is specific to a particular infection. (B) Unsupervised clustering of parasites by ADMIXTURE, each bar denotes the proportion of ancestry a parasite derives from 31 latent populations. (C) Mitochondrial haplotypes for each parasite. Red blocks denote the reference allele, blue the alternative and orange where mixed base calls were observed. The ADMIXTURE proportions and mitochondrial haplotypes are oriented to be below the branch tip of the same parasite. (D) The location of cells from each infection subject to single-cell sequencing in the upper panel.

We used clustering estimated by the UPGMA tree, ADMIXTURE and mitochondrial genotypes (Fig. 3c) to distinguish between infections where diversity results from superinfection or coinfection. Mitochondrial genome sequences are useful markers in this context as they are uniparentally inherited in malaria infections and do not undergo recombination. However, there are few confidently scored mutations in our mitochondrial genome data (n=10) limiting the resolution of our inference. To classify potential superinfections we first identified infections which contained putatively unrelated parasites. Based upon the UPGMA tree 6 infections (MAL5, MAL15, MAL17, MAL23, MAL24, MAL48, Fig. 3d) contained sequences that cluster more closely to other infections than to sequences from the same infection. For 3 of these infections (MAL5, MAL23, MAL24) ADMIXTURE results showed clustering which was congruent with the UPGMA tree. For example, in MAL5 there were 2 distinct clusters by both ADMIXTURE and the UPGMA tree which were in agreement and both of these clusters had a unique mitochondrial genotype. The remaining 3 infections (MAL15, MAL17, MAL48) each showed discordant clustering between the methods. For instance MAL15 shares both ADMIXTURE clusters and mitochondrial genotypes between parasites which were separated by the tree. Based upon this analysis MAL5, MAL23 and MAL24 show evidence of superinfection, while the diversity present MAL15, MAL17 and MAL48 can be explained by co-transmission alone.

## Recent ancestry of individual infections

To better characterize levels of relatedness within infections we identified blocks of chromosomes shared identical-by-descent (IBD) between all paired sequences using a hidden Markov model^27^. IBD sharing between clonal bulk sequenced infections was rare, with a mean of 0.73 blocks shared between infections (range 0-5), encompassing a mean of 88.5kb (range 3.8-342.7kb) of each genome, with a mean block length of 50.8kb (range 3.8-142.4kb). In contrast, within infections parasites shared a mean of 13.0 (range 0-30) IBD blocks between parasite genomes, encompassing a mean of 16,334.2kb (range 3.1-20,577.0kb) of each genome, with a mean shared block length of 1,143.6kb (range 3.1-1469.8kb). As we limit inference of IBD to the ‘core’ genome^28^ identical parasites share 20,577kb of their genomes IBD in 14 blocks (one per chromosome). The presence of IBD sharing between individuals supports recent shared ancestry. For 10/15 of the infections there was at least one block of IBD shared in all pairwise comparisons. As our filtering of IBD blocks was limited to >2.5cM we are limited to inference of relatedness over the last 25 generations (~6 years)^29^.

Recent studies have highlighted the power of IBD networks to capture the structure of a parasite population^29^. We built a network of pairwise shared IBD, creating links between parasites with >15% of their genomes shared IBD (Fig. 4, Supplementary File 3). This revealed close connectivity between parasites from the same infection, with much sparser connectivity between parasites from different infections. We observed subdivision within individual infections, supporting many of the observations in Fig. 2. Varying the minimum IBD required to connect genomes allowed us to visualize how relatedness subdivides individual infections (Supplementary File 3). For instance MAL5 and MAL24 where two clusters of parasites were only connected by the sequence derived from bulk sequencing to one another.

**Figure 4.**
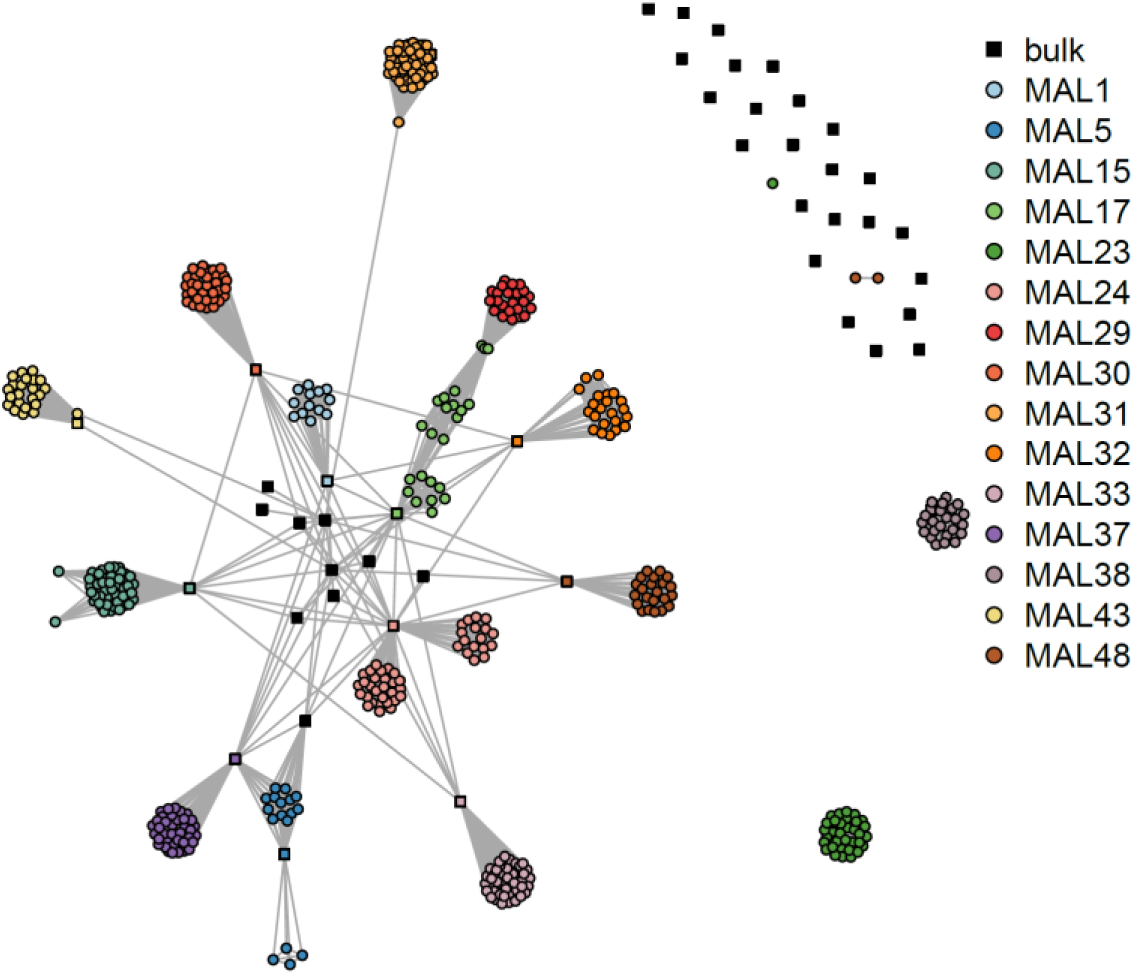
A network representation of pairwise IBD sharing across the genomes. Each node represents a single parasite colored by the infections of origin. Nodes are joined parasites if >15% of the genomes were shared IBD. Each node is colored by the infection it was derived from, with bulk sequences denoted by a square and single cell sequences by a circle.

The distribution of total pairwise shared IBD and the average shared block lengths can be used to infer the relationships between individual genomes^30,31^. We inferred the degree of relatedness from our data using the Estimation of Recent Shared Ancestry (ERSA) algorithm. ERSA estimates relatedness between individuals from distribution of IBD tract lengths (Fig. 5a,b) using assumed unrelated individuals from the same population as a reference. We see a spectrum of relationships within each infection (Fig. 5c, Supplementary File 2). In MAL5 this confirmed the lack of relatedness between the two clusters of parasites, suggesting this infection was the result of a genuine superinfection. However, no other infections can be classified so simply, commonly showing relationships as distant as 4th degree (equivalent to ‘first cousins’). Within our data this suggests that it is not uncommon for parasites to be transmitted through two generations (human-mosquito-human-mosquito-human), with up to four generations of co-transmission seen in our data in infection MAL24 and MAL17. In our data we see only a single unambiguous instance of superinfection of two unrelated parasites with no concurrent co-transmission (MAL5). Across the analysis, the genetic diversity of three infections (MAL17, MAL24 and MAL48) appears to be driven by both superinfection of unrelated parasites and co-transmission of related parasites (in addition to MAL5 where only superinfection is suspected). Surprisingly, we see substantial genetic variation maintained amongst co-transmission parasites. In each of the three infections there were more polymorphic sites segregating between co-infecting parasites than separating superinfecting lineages (Supplementary Data Fig. 6).

**Figure 5.**
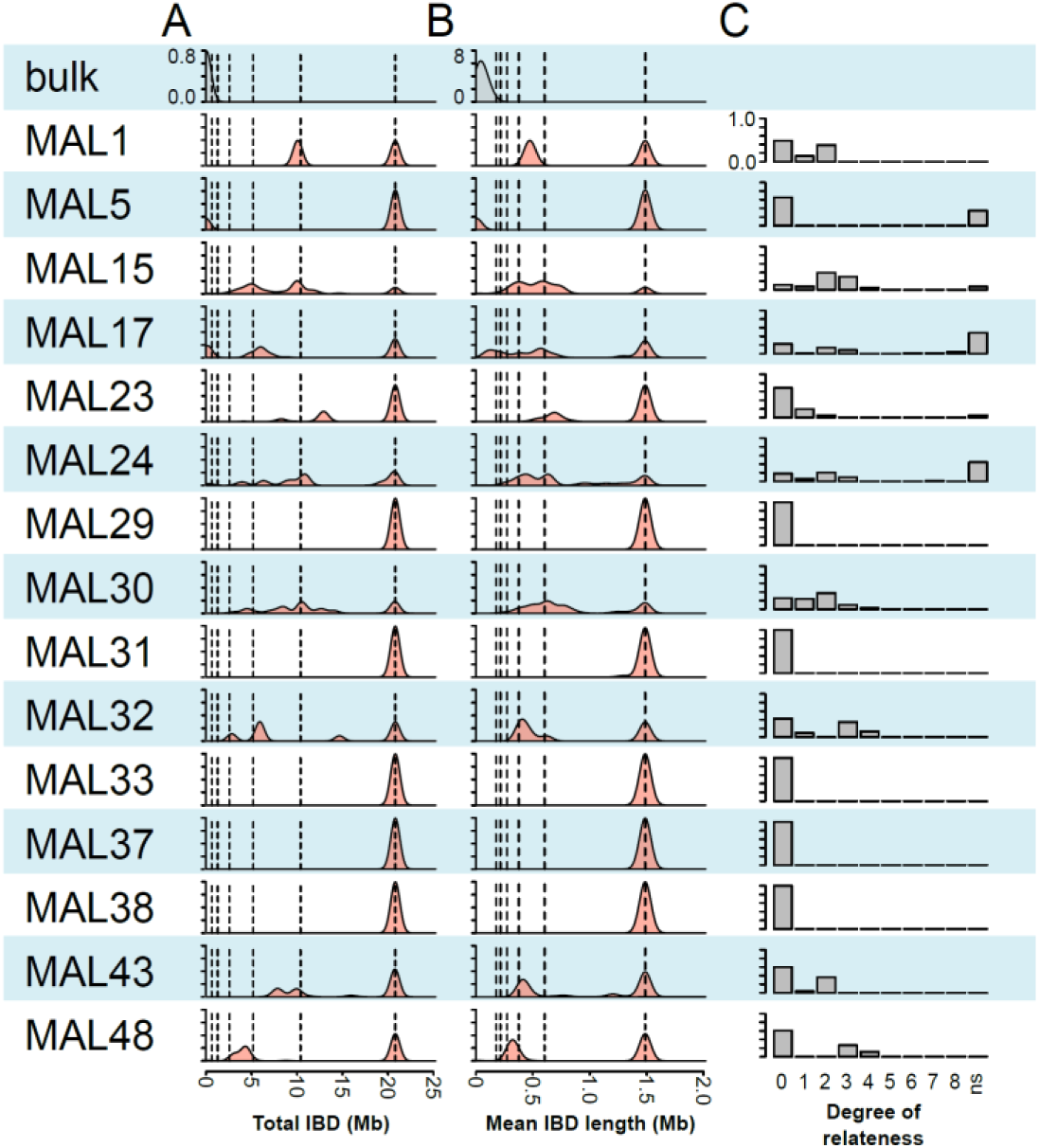
Recent ancestry inferred from IBD sharing. (A) Density plot of the total IBD shared between parasites from a single infection (labeled to the left of the plot). (B) Density plot of the mean IBD block length between parasites from a single infection. (C) Relative frequency of different degrees of relatedness inferred between parasites from the same infection (ns - no significant relatedness observed).

## Discussion

There has been a concerted effort to understand the complexity of malaria infections from either deep sequencing data^13,32,33^, or from genotyping a limited number of markers^12,34^. We show here there is considerable depth to complex infections which may be challenging to infer from bulk analysis alone. Through a combination of deep sequencing of bulk infections and single-cell sequencing we have generated the most comprehensive study of the within-host diversity of malaria infections to date. This provides a much needed standard for developing novel tools for probing the complexity of infections from deep sequencing data. By using multiple estimates of relatedness targeting distinct features of the data we argue that most complex infections result from parasites co-transmitted from single mosquito bites in our dataset. Strikingly, our analysis supports only a single infection where simple superinfection of two unrelated strains has occurred (MAL5), and a further three infections where both superinfection and co-transmission have concurrently contributed to diversity (MAL17, MAL24, MAL48). The remaining infections were either monomorphic or showed strong support for co-transmission of related strains only. In the two infections where we were unable to capture the minor strains (MAL33 and MAL37) patterns of unfixed SNPs within the infection suggest the uncaptured strain was related to the captured strain (Supplementary File 1).

Only parasites which transmit gametes to the same mosquito can produce recombinant offspring. Estimates of the parasite diversity and relatedness within individual mosquitos^35^ (albeit in a distinct population) are in general agreement with our data – most mating is between related parasites. The mechanisms underlying why inbreeding is common, even in high transmission settings, is less clear. Malaria transmission is intense in Chikhwawa^17^ and we expected superinfection to be more prevalent than we observed. A mechanism controlling the outcome of superinfection, perhaps by hepcidin based inhibition of liver development in superinfecting sprozoiites^36^, could explain why we do not see more superinfection. Alternatively, the low numbers of superinfecting parasites emerging from the liver relative to those present in established infections (which may contain 10^11-12^ blood stage parasites) may limit establishment of superinfections. Analysis of parasite diversity is generally limited to single blood draws due to a need to treat symptomatic patients expediently. As this sampling strategy may overlook sub-clones circulating at lower frequencies there may be additional genetic variation which escapes routine analysis.

The depletion of genetic variation during repeated rounds of co-transmission has been previously modelled^6^, suggesting a substantial decline in the number of clones and an increase in average relatedness can arise through a single transmission cycle. Our data suggest that few complex infections have parasites which have been co-transmitted longer than two transmission cycles. We observe substantial genetic variation is maintained despite the bottleneck of mosquito transmission (Supplementary Data Fig. 6) with up to 17 unique haplotypes likely inoculated by a single mosquito. Understanding how patterns of transmission and within host dynamics contribute to the diversity and relatedness structure within malaria infections will be critical to ongoing elimination and control efforts.

## Acknowledgements

We thank all children who participated in this study in Chikhwawa, Malawi. We also thank Andrew Mtande, Ruth Daiman and Miriam Phiri for their help with participant recruitment and clinical management. We are also grateful to Clement Masesa and Lumbani Makhaza for designing our data collection tool and for managing the study database. This study was supported by a Wellcome Trust Intermediate Fellowship in Tropical Medicine and Public Health (Grant # 099992/Z/12/Z to SCN) and an NIH grant (NIAID AI110941-01A1 to IHC). FACS data were generated in the Flow Cytometry Shared Resource Facility (supported by UTHSCSA, NIH-NCI P30 CA054174, UL1RR025767 (CTSA)).

## Author Contributions

S.C.N., S.G.T., R.S.H, S.A.W., D.J.T, T.J.C.A and I.H.C designed the study. S.G.T. and I.H.C. developed tools. S.C.N., S.G.T., K.G., S.N., A.D., C.J., R.G., B.D., and I.H.C. performed experiments. S.C.N., S.K., and D.J.T. collected samples. S.C.N, S.G.T., T.J.C.A and I.H.C. wrote the paper.

## Author Information

The authors declare no competing interests. Correspondence and requests for materials should be addressed to. Raw sequence data has been deposited at the sequence read archive (https://www.ncbi.nlm.nih.gov/sra) under study number SRP155167.

## Methods

### Sample Collection

Malaria-infected blood samples (5 ml; thin smear parasitaemia: 0.2 to 21.8%) were obtained prior to treatment from children aged 19 to 116 months old presenting to Chikhwawa District Hospital in Malawi with uncomplicated *P. falciparum* malaria from February to July 2016. Blood samples were collected in Acid Citrate Dextrose tubes (BD, UK) following consent from parents or guardians, and transported in an ice-cold container to our laboratory in Blantyre, Malawi, for processing. Half of each blood sample was washed using incomplete RPMI 1640 media (Sigma-Aldrich, UK) and cryopreserved in glycerolyte 57 solution (Fenwal, Lake Zurich, IL, USA). Parasites used in fluorescence-activated single-cell sorting were cultured from this sample. The second half of the sample was filtered using CF11 columns to deplete human leucocytes^37^, and was stored at −80°C until needed. Parasite DNA was extracted from this sample using a DNA Mini Kit (QIAGEN, USA) and directly sequenced on an Illumina HiSeq instrument. Ethical approval for this study was obtained from the University of Malawi College of Medicine and Ethics Committee (Protocol number P.02/13/1528) and the Liverpool School of Tropical Medicine Research Ethics Committee (Protocol number 14.035).

### Cell culture and FACS sorting

Approximately 1 mL of frozen sample was thawed at 37°C and parasites were revived (~200ul recovered pellet, ~1% parasitemia). Half of the recovered sample was frozen for bulk DNA extraction and analysis. The other half was grown in 8 mL complete media for 40 hours to allow for parasite progression to late stages, which generates higher quality genomic data after MDA and library preparation^11^. ~8 ul of infected red blood cell pellet was stained in 10 mL PBS which included 5 ul of Vibrant DyeCycle Green at 37C with intermittent mixing for 30 minutes. Cells were washed once in PBS and individually sorted by FACS, gating for trophozoite and schizont-stage parasites.

### Single-cell Sequencing

Library preparation for individually sorted late-stage parasites was carried out using the Qiagen Single-Cell FX DNA kit without library amplification according to manufacturer’s instructions. Library products were analyzed by TapeStation and included off-target peaks typical of MDA DNA inputs. Adapter-ligated DNA products were quantified by KAPA Hyperplus Kits. All sequencing was performed on an Illumina HiSeq 2500.

### Sequence analysis

Median read depth of WGA single-cells was 28.3 (interquartile range (IQR) 12.5-46.4) with median of 90.5% (IQR 78.1-96.0%) of the genome covered by at least one read. In contrast the non-WGA samples had a median read depth of 31.11 (IQR 20.93-48.37) and a median of 95.8% (IQR 93.1-97.4%) of the genome covered by at least one read. A potential source of error in single-cell genomics is the inclusion of exogenous DNA amplified alongside the target genome in downstream analysis. As an initial indication of the potential of non-target DNA being introduced to our analysis we first examined the proportion of reads mapping to the *P. falciparum* genome^24^ in each sequence. We observed a median of 93.3% (IQR 87.0-95.4%) of reads map to the parasite genome for single-cell sequences, compared to 35.7% (IQR 19.7-48.5%) for bulk patient samples and 79.4% (IQR 74.5-86.9%) for clonally expanded samples suggesting our stringent handling protocols were effective at eliminating environmental DNA. For a more rigorous test we identified lines with potential cross contamination based on unfixed basecall frequency. As the parasite genome is haploid during blood stages all variants are expected to be fixed in genome sequencing data. The highly AT-rich and repetitive nature of the parasite genome makes alignment challenging, generating false positive unfixed variants in clonal lines. After excluding highly error-prone genomic regions (calls outside of the “core genome”^28^ or within microsatellites) we measured the proportion of mixed base calls (>5% of reads at a locus mapping to the minority allele) at high confidence biallelic SNPs (>10 reads mapped, VQSLOD>0, GQ>70). Using the cloned lines and bulk population samples as a guide we estimated 1% as an appropriate threshold for excluding putatively mixed lines (Supplementary Data Fig. 1).

### Estimating the complexity and diversity of bulk sequenced samples

F_WS_ was calculated in moimix (https://github.com/bahlolab/moimix) for all bulk patient samples. We estimated the number of unique haplotypes and their sequence from deep sequence of bulk infections using DEploid^13^ v0.5 (https://github.com/mcveanlab/DEploid). We used 10,997 HQ SNPs with a MAF >5%. For a reference panel we used 10 bulk Malawian samples presumed to be clonal (F_WS_>0.95) and population level allele frequencies from across the complete bulk sequencing data. We inferred the most likely number of haplotypes (K) using the command:./dEploid -ref sample_reference_allele_counts.txt -alt sample_alternative_allele_counts.txt -plaf population_allele_freq.txt -o sample_out -ibd -noPanel -exclude highly_variable_sites.txt -sigma 7 -seed 2

### Estimating relatedness between sequences

SNP data were imported into R using SeqArray^38^. Between all samples passing quality control we calculated the proportion of shared alleles and using SNPs which were at >5% MAF in the bulk sequenced samples. We used a distance matrix generated from this data (1-pairwise allele sharing) to build a UPGMA tree (Fig. 2). We also used this statistic to estimate the number of unique haplotypes in each infection by collapsing together sequences which differed at <1% of sites. Rarefaction of haplotype abundance was performed using the rareNMtests package^39^ in R. We performed unsupervised clustering of the sequence data using ADMIXTURE v1.3^26^ (https://www.genetics.ucla.edu/software/admixture/). This again used sites with a MAF of >5% across the bulk sequenced data. We clustered data using K of 2-40 seeing a minima of CV error at K=31 (Supplementary Data Fig. 5). We called regions of IBD between all samples passing quality control using hmmIBD v2.0.0^27^ (https://github.com/glipsnort/hmmIBD). We performed maximum-likelihood estimation of recent shared ancestry using ERSA 2.0^30,31^ (http://www.hufflab.org/software/ersa/) using the output from hmmIBD using the flags -- min_cm=1.5 --adjust_pop_dist=true --number_of_chromosomes=14 --rec_per_meioses=19. We converted the basepair positions to a uniform genetic map using the scaling factor 1cM=9.6kb^40^ and excluded IBD chunks <1cM in length. As identical clones are not specifically modelled in ERSA we excluded these from analysis, though their abundance is shown in the ‘0’ bar in Fig. 3c. All other statistical analysis and visualization was performed in R v3.4.0^41^.

